# Inosculation as a tool for living architecture: Methods and early results

**DOI:** 10.1101/2024.10.30.621064

**Authors:** P. Faucon, A. Fertet, A. Bombail, C. Gouguec, C. Montil, A. de la Sayette

## Abstract

Among the ways to introduce more plants within the city, living architecture could lower material required for building a house. However, trees are not supposed to have a house-like shape, so it is needed to give them this shape. That is why, we choose to investigate the assembly of living trees through inosculation. To do so, we initiated this work in March 2021, in our greenhouse, on five species: *Acer platanoides, Celtis australis, Cornus Mas, Corylus colurna* and *Pyrus communis*. We tested two types of contact (parallel or perpendicular) between two trees from the same species with scraped bark, or not, maintained by diverse tools (screw, fishing line, buddy tape and flexible stretch tie). From this, we get interesting anastomosis results with half of the trees showing signs of inosculation by spring 2022.

## INTRODUCTION

According to the Intergovernmental Panel on Climate Change (IPCC), future changes to both climate and urbanization will enhance warming in cities (Shukla et al., 2019). This effect is expected to be increased during heat waves, making cities very unpleasant for people. Moreover, ever-growing cities induce a bad air quality and so premature death. Climate change is also expected to intensify rainfall intensity (Stocker et al., 2013) which in addition to soil sealing is already at the root of flooding. Still according to the IPCC, the implementation of urban green infrastructure can contribute to climate change mitigation and adaptation (Shukla et al., 2019). Consequently, it is necessary to increase the place of the plants in the city. For now, in city, plants are limited to parks, green roofs and street decorations, but this could change in the upcoming years with the development of living architecture.

Living architecture is defined as the use of natural living organisms to build sustainable construction (Vallas and Courard, 2017). Such a strategy of using living trees as the structure is not new as it has been used for centuries. Indeed, bridges made of living roots of *Ficus elastica* can be found in the Indian State Meghalaya with some of them as old as several hundreds of years (Ludwig et al., 2019). Such bridges are made by guiding roots from one shore to the other across the bamboo deadwood framework (Ludwig et al., 2019). Other examples of living bridges can be found in Japan across the Iya Valley and are made of *Wisteria floribunda* (Vallas and Courard, 2017). In both cases, shaping the plant was limited to reaching the other shore. However, it is possible to go further and shape trees to required forms. Such idea can be found in the work of John Krubsack (1858-1941), who realized the first living-chair made in 11 years from 32 seedlings, and Axel Erlandson (1884-1964), who built more than 70 tree sculptures (Vallas and Courard, 2017). Nowadays, such works can be found in Germany with the Sanfte Strukturen team, who built a palace made of 10’000 bend *Salix alba*, and with the Ludwig group from the Technical University of Munich. In 2005, Prof. Dr.-Ing. Ferdinand Ludwig started to develop a new concept in architecture which he called “Baubotanik”. “Baubotanik” consists of building a conventional structure to guide trees to the desired form and once the shape is achieved, to remove non-living parts of the building (Ludwig et al., 2012). By doing so, the Ludwig group built successfully a first 8 meters high tower with *Platanus acerifolia* in 2009 and transposed it to a larger one in 2012 in Nagold, Germany (Ludwig, 2014).

One of the keys of living architecture is to maintain the shape of the tree and so the shape of the structure. To do so, it is necessary to connect plants so that they merge into a single physiological unit with mechanically strong junctions (Ludwig et al., 2012). Such a method is called inosculation, or grafting, and was invented approximately 2,000 years before the Common Era (Mudge et al., 2009). Indeed, in horticulture, grafting is used for the multiplication of fruit trees, vine or certain vegetables under a greenhouse. However, in horticulture, the method consists of adding on a plant with roots, called rootstock, a part of another plant, or scion, while in living architecture, it consists of making from two plants one, at the trunk level, while keeping their integrity. In both cases, grafting means deliberate fusion of plant parts so that vascular continuity is established between them (Gaut et al., 2019). From a cellular point of view, the vascular cambium, which is the main grown tissue and contains meristematic cells integral to repair, is the one involved in grafting. The cambium is surrounded by the sapwood, secondary xylem, and the live bark, secondary phloem. To be successful, grafting required three physiological steps (Yin et al., 2012; Wang et al., 2017; Gaut et al., 2019): 1) First, parts (trunks or scion and rootstock) must adhere which typically occurs through the excretion of pectin; 2)Second, callus develops from both parts; 3) Third, callus and cambium differentiate into xylem and phloem strands establishing the vascular connection between the two parts and allowing the long-distance movement of water and solutes. Such a technique is mastered by the Ludwig Group for *S. alba* and *P. acerifolia* by making a perpendicular contact between trunks maintained by a screw (Ludwig et al., 2008, 2012; Ludwig, 2014; Middleton et al., 2022).

In order to extend available tree species for living architecture, in our work, we aimed to study grafting to five new species: *Acer pseudoplantanus, Celtis australis, Cornus Mas, Corylus colurna* and *Pyrus communis*; all of them suitable for grafting thanks to their thin bark or growing in urban dry condition. To do so, we studied two types of contact: parallel or perpendicular; between two trees from the same species, with scratched bark or not. We also studied different tools: screw, fishing line, grafting tape and flexible stretch tie; to maintain contact between the trees. We initiated this work in march 2021, in our greenhouse in Rochefort (Charente-Maritime, France) and we observed promising results from spring 2022. Indeed, on the 60 tests that we set, half of them showed early signs of inosculation.

## MATERIALS AND METHODS

### Plant material and supplies

8 *A. pseudoplantanus*, 16 *C. australis*, 8 *C. Mas*, 8 *C. colurna* and 16 *P. communis* were purchased from *PÉPINIÈRES DE CORME-ROYAL* (Corme-Royal, Charente-Maritime, France), as saplings 200/250 cm excepted for 4 *C. australis* ordered as stem trees 8/10. For their growth, we purchased 27 stainless steel 240×40×77 cm vats from *VIDAXL* (Venlo, Limburg, THE NETHERLAND) that we fill with Hydrop® substrate from *PREMIER TECH* (Rivière-du-Loup, Québec, CANADA). We use as fertilizer Osmocote Pro® 8-9 months (18-9-10 + 2 MgO + OE) from *ICL* (Tel Aviv, ISRAEL). The trees were grown on Hydrop® substrate under our glace greenhouse without artificial light. The substrate was supplied every six months with 300 g of fertilizer for 1m3 of substrate. To maintain contact between the trunks, we used a fishing line Spiderwire Ultracast Ultimate Braid 8H 110 m Green 0.250 mm (*PURE FISHING*, Columbia, South Carolina, USA), a grafting tape (*LONODIS*, Saint Chef, Isère, FRANCE), a flexible stretch tie Scoubitol® (*TOLTEX*, Argenton-les-Vallées, Deux-Sèvres, FRANCE) and/or a wood screw of stainless steel 3×29 mm.

### Grafting procedures

The trees were grown for 2 months prior to grafting, from February 2021 to April 2021. Then tree pairs have been manually joined according to two types of contact: perpendicular or parallel (**Figure 1**). Perpendicular contact consists of the simple juxtaposition of the two trunks with an angle comprised between 90° and 30°. Parallel contact required to bend trunks to get a flat surface of contact about 30 cm. Because the study took place under a greenhouse, prior to that contact, the bark of some pairs of trees was scratched bark to mimic the natural scrap induced by the wind in natural conditions. To do so, we remove the first layer (1 mm) of bark with a scalpel. Based on literature and experiences from our team, 4 materials was selected to maintain the contact on the long run: fishing line or flexible stretch tie, as thin links, grafting tape, commonly used in multiplication, and screw, used by the Ludwig group. Only the grafting tape and the flexible stretch tie were used alone for some pairs, all other pairs showed two or three linking materials (for complete data, see **Supplementary S1**).

**Figure 1.**
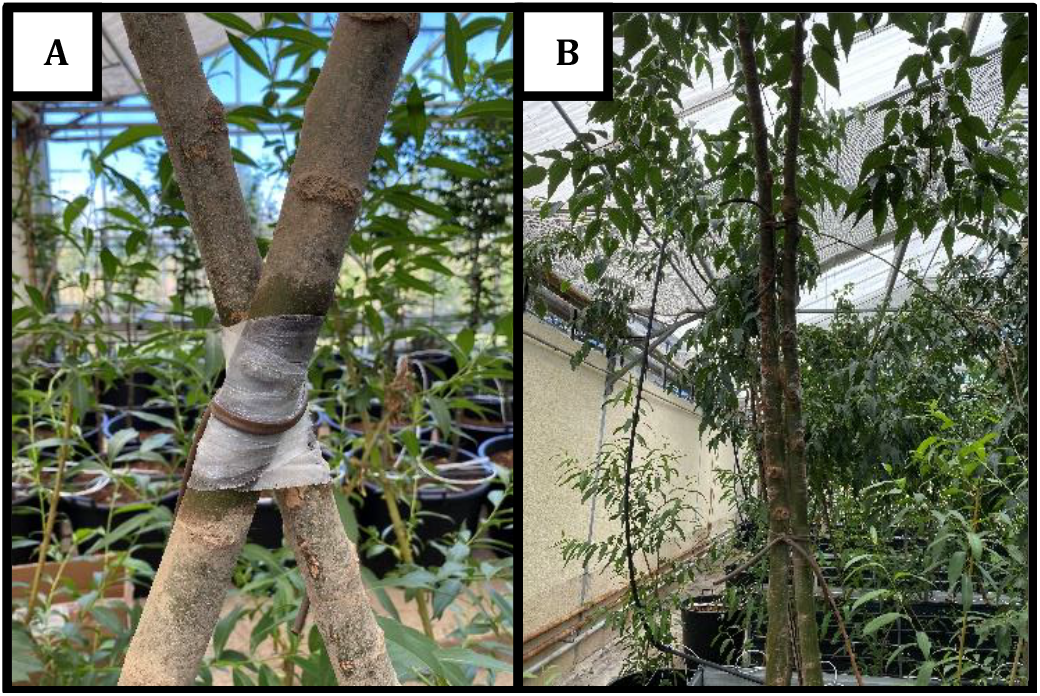
The two types of contact. **A**. A pair of Celtis australis in a perpendicular contact maintained by grafting tape and a flexible stretch tie. **B**. A pair of Pyrus communis in a parallel contact maintained by two flexible stretch ties.

### Non-destructive evaluation of inosculation

This experiment is expected to run over a few years. Consequently, the observation of early signs of inosculation was made one year after grafting, in march 2022, by looking at the contact area without removing linking materials. To do so, callus or early sign of callus have been looked for and if such a structure was visible the grafting was considered as well underway. For some pairs, because of linking materials, it was impossible to determine whenever inosculation started or not (**Figure 2**).

**Figure 2.**
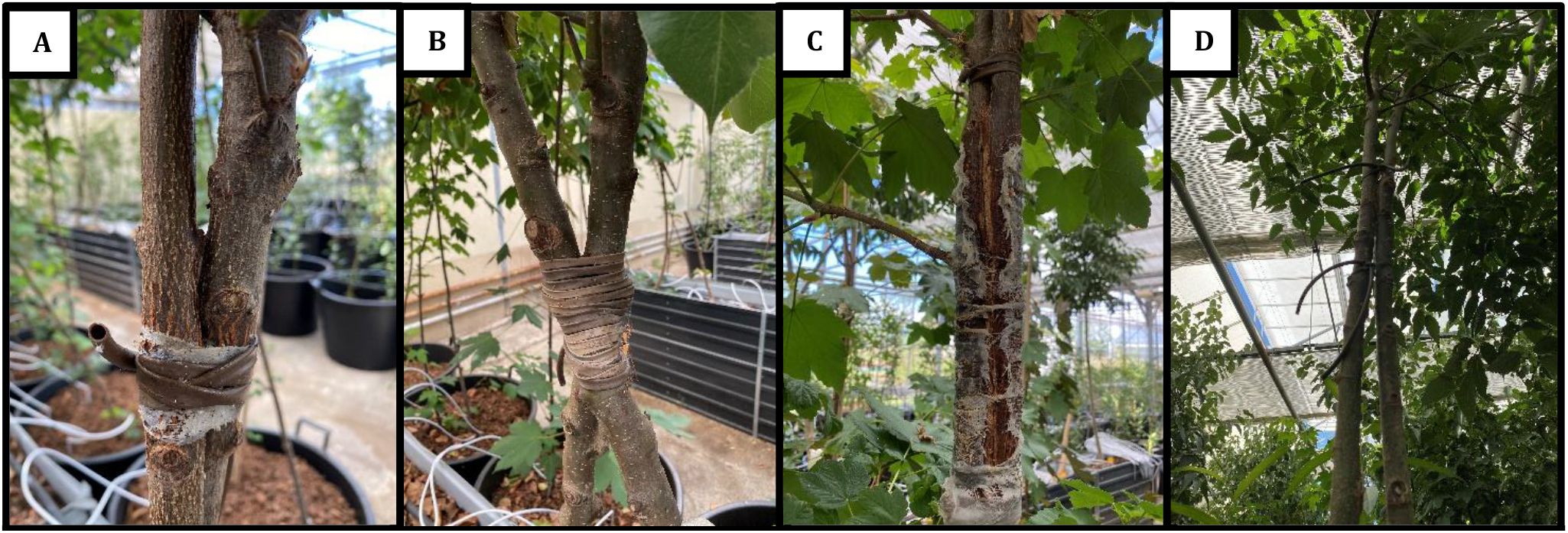
Non-destructive evaluation of inosculation on (A and B) perpendicular contact and (C and D) parallel contact. **A**. Perpendicular contact with early signs of inosculation. **B**. Early sign of inosculation impossible to observe due to linking material. **C**. Parallel with clear early signs of inosculation. **D**. No sign of inosculation.

## RESULTS AND DISCUSSION

### Influence of contact type on grafting

Success rate calculated according to the type of contact shows differences according to the species (**Table 1**): Parallel contact seems to work better with *Cornus mas* because the two pairs tested in that condition showed signs of inosculation. *Acer pseudoplatanus* gave the best results with the same condition. *Celtis australis* planted in saplings shows also good results in the parallel type of contact with a success rate of 73 %, but seems to be interesting in the perpendicular contact with 40 % of success. However, stem trees did not show any signs of inosculation. *Corylus colurna* is, at that time, the most adaptive of the five species to inosculation with a high success rate (75 %) regardless of the approach method. Finally, *Pyrus communis* follows the pattern of other species by showing better results when the type of contact is parallel between the two trunks with 80 % compared to 36 % for perpendicular one.

**Table 1.** Results of early signs of inosculation according to type of contact.

| Species | Type of contact | Number of pairs concerned | Pairs showing signs of inosculation | Ratio of success |
| --- | --- | --- | --- | --- |
| *Acer pseudoplatanus* | Parallel | 4 | 3 | 75% |
|  | Perpendicular | 4 | 0 | 0% |
| *Celtis australis* sapling | Parallel | 11 | 8 | 73% |
|  | Perpendicular | 5 | 2 | 40% |
| *Celtis australis* stem tree | Perpendicular | 4 | 0 | 0% |
| *Cornus mas* | Parallel | 2 | 1 | 50% |
|  | Perpendicular | 6 | 1 | 17% |
| *Corylus colurna* | Parallel | 4 | 3 | 75% |
|  | Perpendicular | 4 | 3 | 75% |
| *Pyrus communis* | Parallel | 5 | 4 | 80% |
|  | Perpendicular | 11 | 4 | 36% |
| **Overall** | Parallel | 26 | 19 | 73% |
|  | Perpendicular | 34 | 10 | 29% |

Regarding these results, we believe that our non-destructive method to evaluate early signs of grafting may induce a bias with respect to the type of contact. Indeed, the area of contact in parallel modality is larger than the one of perpendicular contacts. Meaning that there is greater opportunity to observe signs of inosculation in the parallel type of contact than in the perpendicular one, which we observed with mean success rates of 73 % against 29 %, respectively. When looking at the species level, we observed two interesting results that might need further investigation. First, the fact that we did not observe any early signs of grafting in *C. australis* could be due to the late planting in bare roots which is undoubtedly more expensive in energy than the younger subjects. Their low growth and the age of the trees undoubtedly influences the success or the speed of the anastomosis. For now, we cannot exclude that the stem trees will merge at some point, it is possible that it takes more time. Then, it could be interesting to investigate the reasons behind the high rate of success observed in *C. colurna*. For now, we hypothesize that it could be linked to its squamous bark or that we think.

### Influence of bark integrity on grafting

In the wild, the wind scratches the bark of trees in contact by scrubbing them against each other. Because our experiment takes place under a glass greenhouse, the wind cannot play that role. To overcome that lack, we scratch the bark of the trees by hand with a scalpel. However, we wanted to study the interest of scraping the bark so on the 60 pairs of trees in our experiment, we scratched the barks of 45 pairs of trees (**Table 2**). Of these 45 pairs of trees, 23 show early signs of inosculation, 51 % of success, with respect to the 53 % of success for pairs with intact bark (8 out of 15). Such a result seems to suggest that there is no effect of scratching the bark prior to contact but when looked through species, the conclusion might be different. Indeed, results for *A. pseudoplantanus* suggests an interesting impact of scratching in the success of inosculation with a ratio of success about 43 % against 0. Same conclusion could be made about *C. mas*, with a ratio of success about 75 % against 25 %. While in *P. communis*, bark integrity doesn’t seem to be the most important factor.

**Table 2.**
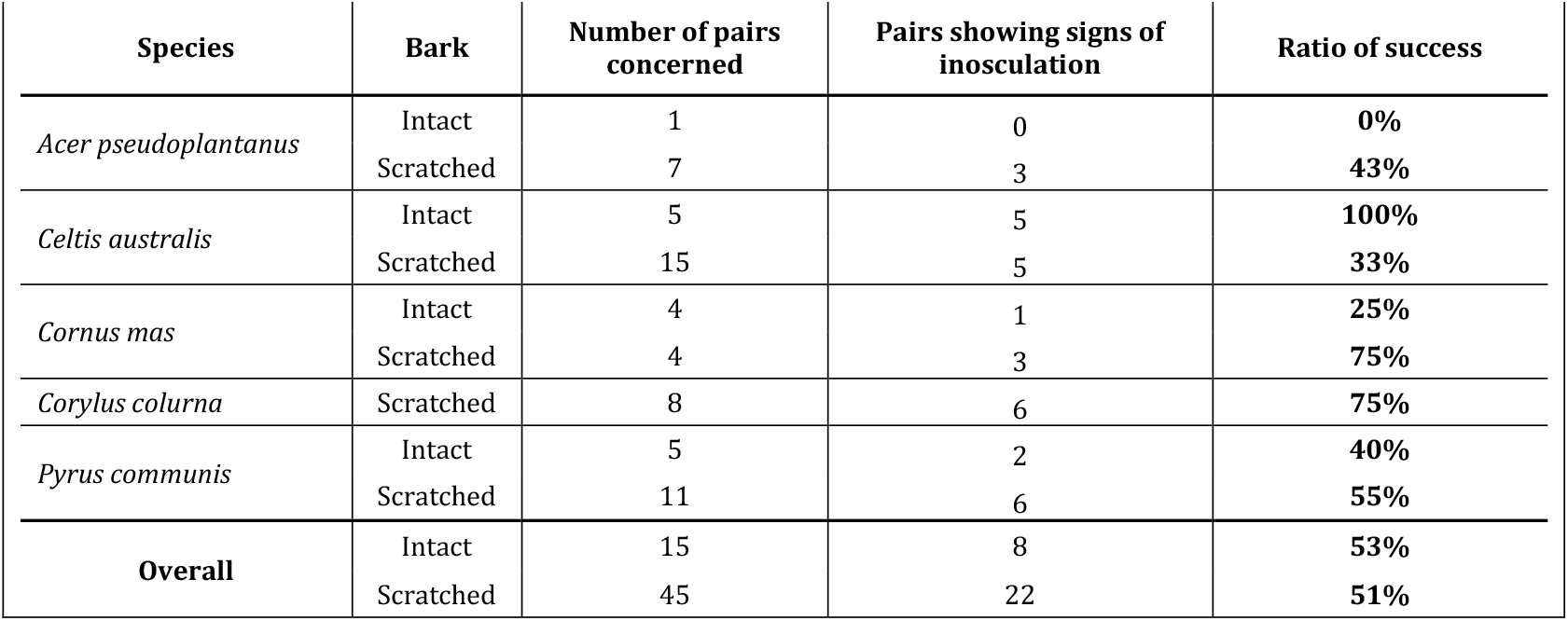
Results of early signs of inosculation according to the state of integrity of the bark.

However, as we saw previously, *C. australis* might be an out layer. Indeed, if we discard data from this species, we found a success rate of 56 % for scratched pairs with respect to the 30 % for non-scratched ones. We agree that this looks like cherry-picking, that is why we cannot conclude in the interest of scratching yet. Furthermore, the population of non-scratched pairs is too low compared to the scratched, which might complicate to conclude on this method.

### Influence of linking materials on grafting

In order to maintain the contact between the two trees of the pairs, we used 4 linking materials: grafting tape, screw, fishing line and a flexible stretch tie. We used these materials mostly in combination of 2 (40 pairs), but also in combination of 3 (6 pairs, tied only with grafting tape, screw and the flexible stretch tie) or alone for 14 pairs, linked with grafting tape only (7) or with the flexible stretch only (4) (for complete data, see **Supplementary S2**).

The overall results (**Table 3**) show that in our condition and with our method of evaluation of early signs of grafting, the linking material with the greatest ratio of success is the fishing line with a ratio of 79 %, 15 out of 19, followed by the grafting tape 53 % and the screw, 45 %. However, the poor ratio of success obtained with the flexible stretch tie seems to indicate that this material is not suitable for such a method of grafting.

**Table 3.** Results of early signs of inosculation according to the linking materials.

|  |  |  |  |  |
| --- | --- | --- | --- | --- |
| Species | Linking material | Number of pairs concerned | Pairs showing signs of inosculation | Ratio of success |
| *Acer pseudoplatanus* | Grafting tape | 8 | 3 | 38% |
|  | Screw | 5 | 2 | 40% |
|  | Fishing line | 2 | 1 | 50% |
|  | Flexible stretch tie | 3 | 0 | 0% |
| *Celtis australis* | Grafting tape | 20 | 10 | 50% |
|  | Screw | 6 | 4 | 67% |
|  | Fishing line | 8 | 6 | 75% |
|  | Flexible stretch tie | 3 | 0 | 0% |
| *Cornus mas* | Grafting tape | 6 | 3 | 50% |
|  | Screw | 3 | 1 | 33% |
|  | Fishing line | 2 | 2 | 100% |
|  | Flexible stretch tie | 3 | 1 | 33% |
| *Corylus colurna* | Grafting tape | 8 | 6 | 75% |
|  | Screw | 4 | 3 | 75% |
|  | Fishing line | 2 | 2 | 100% |
|  | Flexible stretch tie | 1 | 1 | 100% |
| *Pyrus communis* | Grafting tape | 13 | 7 | 54% |
|  | Screw | 5 | 3 | 60% |
|  | Fishing line | 3 | 3 | 100% |
|  | Flexible stretch tie | 4 | 1 | 25% |
| **Overall** | Grafting tape | 55 | 29 | 53% |
|  | Screw | 23 | 13 | 57% |
|  | Fishing line | 17 | 14 | 82% |
|  | Flexible stretch tie | 14 | 3 | 21% |

The grafting tape was applied in a large population, 55 out of 60, because it seems the easiest material to implement and the least harmful for the trunks. However, we have not tested the modality without grafting tape enough to conclude, yet. Concerning the fishing line, pairs with this material showed good results, but it lacerated the bark of *C. mas* and *A. pseudoplatanus* so we removed it after a certain period for these two species. Consequently, this material doesn’t seem relevant to us. The marginal use of the flexible stretch tie does not allow us to conclude on its effectiveness, but the tendency is not in favour of this material. Finally, the screw was probably the tougher material for the trees because it had to go through the entire wood of at least one of the trees of the pair, but we had interesting results, especially in the case of *C. colurna* with 75 % of success.

With respect to the combination of linking material, for the 14 pairs with only 1 material, we get 35 % of success and for the 40 with 2 materials 63 %, while for the 6 pairs with 3 materials, only 1 showed signs of inosculation (meaning 17 %). Consequently, it doesn’t seem that multiplying linking materials over 2, is a good way to induce inosculation. Furthermore, we may have not found the perfect material or combination of materials for each of these species but these preliminary results are promising.

## CONCLUSION

In this study, we presented preliminary results for a work aiming to test inosculation between trees in 5 species: *Acer pseudoplantanus, Celtis australis, Cornus Mas, Corylus colurna* and *Pyrus communis*; according to two types of contact: parallel or perpendicular; with scratched bark or not, by using 4 different tools: screw, fishing line, grafting tape and flexible stretch tie; to maintain contact between the trees. This work was initiated in March 2021 so this study shows the results after one year. That is why these results must be considered as preliminary and we expect inosculation to appear for more pairs in the upcoming year. For now, if we need to conclude on this study, the species with the best ratio of success is *Corylus colurna*, the most appropriate type of contact is the parallel one maintained with grafting tape. However, we consider this study a success as of now.

This study allowed us to practice finding our bearings in a field of research which is a grey area. Indeed, the plant nursery environment suffers from real loss of know-how in anastomosis between two living trees. When we built this study, we planned to let inosculation run over the years and consequently does not allow us to destroy to image or see really what is going on between the trees. That is why we initiated a new study in April 2022 focused on two species with multiple replicates that we will destroy over the year to image the process of inosculation at a macroscopic level.

In the long run, with trees accepting anastomosis with multiple fellows, we could delimit square shapes with living trees that can be the bases of living architecture, and, on the very long run, the bases of living houses. However, such an idea could face reglementary or acceptability issues. Such a strategy could be reasonably used to build living street furniture such as bus shelters or bin storages.

## ACKNOWLEDGEMENTS

We acknowledge support from all members of ARRDHOR CRITT Horticole.

## Supplementary

**Supplementary S1.** Complete data used in this study.

| Supplementary S1. Complete data used in this study |  |  |  |  |  |  |  |
| --- | --- | --- | --- | --- | --- | --- | --- |
| Pair # | Species | Type of contact | Bark | Linking material #1 | Linking material #2 | Linking material #3 | Signs of inosculation |
| 1 | *Acer pseudoplatanus* | Parallel | Scratched | Grafting tape | Fishing line | - | Yes |
| 2 | *Acer pseudoplatanus* | Parallel | Scratched | Grafting tape | Fishing line | - | Yes |
| 3 | *Acer pseudoplatanus* | Parallel | Scratched | Grafting tape | Screw | - | - |
| 4 | *Acer pseudoplatanus* | Parallel | Scratched | Grafting tape | Screw | - | - |
| 5 | *Acer pseudoplatanus* | Parallel | Scratched | Grafting tape | Screw | - | Yes |
| 6 | *Acer pseudoplatanus* | Perpendicular | Intact | Grafting tape | Flexible stretch tie | Screw | - |
| 7 | *Acer pseudoplatanus* | Perpendicular | Scratched | Grafting tape | Flexible stretch tie | - | - |
| 8 | *Acer pseudoplatanus* | Perpendicular | Scratched | Grafting tape | Flexible stretch tie | Screw | - |
| 9 | *Celtis australis* saplings | Parallel | Intact | Grafting tape | Fishing line | - | Yes |
| 10 | *Celtis australis* saplings | Parallel | Intact | Grafting tape | Fishing line | - | Yes |
| 11 | *Celtis australis* saplings | Parallel | Intact | Grafting tape | Fishing line | - | Yes |
| 12 | *Celtis australis* saplings | Parallel | Intact | Grafting tape | Fishing line | - | Yes |
| 13 | *Celtis australis* saplings | Parallel | Scratched | Grafting tape | Fishing line | - | - |
| 14 | *Celtis australis* saplings | Parallel | Scratched | Grafting tape | Fishing line | - | - |
| 15 | *Celtis australis* saplings | Parallel | Scratched | Grafting tape | Fishing line | - | Yes |
| 16 | *Celtis australis* saplings | Parallel | Scratched | Grafting tape | Fishing line | - | Yes |
| 17 | *Celtis australis* saplings | Parallel | Scratched | Grafting tape | Screw | - | - |
| 18 | *Celtis australis* saplings | Parallel | Scratched | Grafting tape | Screw | - | Yes |
| 19 | *Celtis australis* saplings | Parallel | Scratched | Grafting tape | Screw | - | Yes |
| 20 | *Celtis australis* saplings | Perpendicular | Intact | Grafting tape | Screw | - | Yes |
| 21 | *Celtis australis* saplings | Perpendicular | Scratched | Grafting tape | Flexible stretch tie | - | - |
| 22 | *Celtis australis* saplings | Perpendicular | Scratched | Grafting tape | Flexible stretch tie | - | - |
| 23 | *Celtis australis* saplings | Perpendicular | Scratched | Grafting tape | Flexible stretch tie | Screw | - |
| 24 | *Celtis australis* saplings | Perpendicular | Scratched | Grafting tape | Screw | - | Yes |
| 25 | *Celtis australis* stem trees | Perpendicular | Scratched | Grafting tape | Flexible stretch tie |  | - |
| 26 | *Celtis australis* stem trees | Perpendicular | Scratched | Grafting tape | Flexible stretch tie |  | - |
| 27 | *Celtis australis* stem trees | Perpendicular | Scratched | Grafting tape |  |  | - |
| 28 | *Celtis australis* stem trees | Perpendicular | Scratched | Grafting tape |  |  | - |
| 29 | *Cornus mas* | Parallel | Scratched | Grafting tape | Fishing line | - | Yes |
| 30 | *Cornus mas* | Parallel | Scratched | Grafting tape | Fishing line | - | Yes |
| 31 | *Cornus mas* | Perpendicular | Intact | Flexible stretch tie | Screw | - | Yes |
| 32 | *Cornus mas* | Perpendicular | Intact | Grafting tape | Flexible stretch tie | - | - |
| 33 | *Cornus mas* | Perpendicular | Intact | Grafting tape | Screw | - | - |
| 34 | *Cornus mas* | Perpendicular | Intact | Grafting tape | Screw | - | - |
| 35 | *Cornus mas* | Perpendicular | Scratched | Flexible stretch tie | - | - | - |
| 36 | *Cornus mas* | Perpendicular | Scratched | Grafting tape | - | - | Yes |
| 37 | *Corylus colurna* | Parallel | Scratched | Grafting tape | Fishing line | - | Yes |
| 38 | *Corylus colurna* | Parallel | Scratched | Grafting tape | Fishing line | - | Yes |
| 39 | *Corylus colurna* | Parallel | Scratched | Grafting tape | Screw | - | - |
| 40 | *Corylus colurna* | Parallel | Scratched | Grafting tape | Screw | - | Yes |
| 41 | *Corylus colurna* | Perpendicular | Scratched | Grafting tape | - | - | - |
| 42 | *Corylus colurna* | Perpendicular | Scratched | Grafting tape | - | - | Yes |
| 43 | *Corylus colurna* | Perpendicular | Scratched | Grafting tape | Flexible stretch tie | Screw | Yes |
| 44 | *Corylus colurna* | Perpendicular | Scratched | Grafting tape | Screw | - | Yes |
| 45 | *Pyrus communis* | Parallel | Intact | Grafting tape | Fishing line | - | Yes |
| 46 | *Pyrus communis* | Parallel | Intact | Grafting tape | Screw | - | Yes |
| 47 | *Pyrus communis* | Parallel | Scratched | Grafting tape | Fishing line | - | Yes |
| 48 | *Pyrus communis* | Parallel | Scratched | Grafting tape | Fishing line | - | Yes |
| 49 | *Pyrus communis* | Parallel | Scratched | Grafting tape | Screw | - | - |
| 50 | *Pyrus communis* | Perpendicular | Intact | Flexible stretch tie | - | - | - |
| 51 | *Pyrus communis* | Perpendicular | Intact | Grafting tape | - | - | - |
| 52 | *Pyrus communis* | Perpendicular | Intact | Grafting tape | Screw | - | - |
| 53 | *Pyrus communis* | Perpendicular | Scratched | Flexible stretch tie | - | - | - |
| 60 | *Pyrus communis* | Perpendicular | Scratched | Flexible stretch tie | - | - | Yes |
| 54 | *Pyrus communis* | Perpendicular | Scratched | Grafting tape | - | - | - |
| 55 | *Pyrus communis* | Perpendicular | Scratched | Grafting tape | - | - | - |
| 56 | *Pyrus communis* | Perpendicular | Scratched | Grafting tape | - | - | Yes |
| 57 | *Pyrus communis* | Perpendicular | Scratched | Grafting tape | Flexible stretch tie | - | - |
| 58 | *Pyrus communis* | Perpendicular | Scratched | Grafting tape | Screw | - | Yes |
| 59 | *Pyrus communis* | Perpendicular | Scratched | Grafting tape | Screw | - | Yes |

